# LTB_4_ is converted into a potent human neutrophil NADPH oxidase activator via a receptor transactivation mechanism in which the BLT_1_ receptor activates the free fatty acid receptor 2

**DOI:** 10.1101/2024.11.12.623248

**Authors:** Yanling Wu, Claes Dahlgren, Huamei Forsman, Martina Sundqvist

## Abstract

The endogenous neutrophil chemoattractant leukotriene B_4_ (LTB_4_) is a biased signalling agonist that potently increases the intracellular concentration of free calcium ions ([Ca^2+^]_i_), but alone is a weak activator of the neutrophil superoxide anion (O_2_^-^)-generating NADPH oxidase. However, in this study we show that an allosteric modulator of the free fatty acid 2 receptor (FFA2R) converts LTB_4_ into a potent NADPH oxidase activating agonist. While an allosteric modulation of FFA2R was required for LTB_4_ receptor 1 (BLT_1_R)-mediated activation of the NADPH oxidase, the LTB_4_-induced increase in [Ca^2+^]_i_ was not affected by the modulator. Thus, the biased BLT_1_R signalling pattern was altered in the presence of the allosteric FFA2R modulator, being biased with a preference towards the signals that activate the NADPH oxidase relative to an increase in [Ca^2+^]_i_. Both BLT_1_R and FFA2R belong to the family of G protein-coupled receptors (GPCRs), and our results show that a communication between the activated BLT_1_R and the allosterically modulated FFA2Rs generates signals that induce NADPH oxidase activity. This is consistent with a previously described receptor transactivation (crosstalk) model whereby the function of one neutrophil GPCR can be regulated by receptor downstream signals generated by another GPCR. Furthermore, the finding that an allosteric FFA2R modulator sensitises not only the response induced by orthosteric FFA2R agonists but also the response induced by LTB_4_, violates the receptor restriction properties that normally define the selectivity of allosteric GPCR modulators.

## 1. Introduction

Neutrophils, the most abundant leukocytes in human blood, are of primary importance for our host defence and the fine-tuning/regulation of inflammatory responses [1]. The functions of these cells are regulated by a large number of receptors, many of which belong to the family of G protein-coupled receptors (GPCRs). This is the largest family of human plasma membrane receptors (> 800 members) and is involved in the regulation of a variety of important physiological processes, including inflammation [2, 3]. Accordingly, many available drugs act through GPCRs, and ligands targeting GPCRs are widely being investigated as novel drug candidates for various diseases [4-6]. In general, signalling initiated by agonist binding to GPCRs is based on the activation of intracellular heterotrimeric G proteins [3, 7].The arachidonic acid (AA)-derived dihydroxy acid leukotriene B4 (LTB_4_) is a lipid inflammatory mediator recognised by receptors belonging to the GPCR family [8-10]). Two cognate GPCRs (leukotriene B_4_ receptors [BLTRs]) have been identified for LTB_4_. These BLTRs (BLT_1_R and BLT_2_R) differ in their functional profile and expression pattern as well as in their specificity and affinity for LTB_4_. While BLT_1_R has been suggested to promote pro-inflammatory responses, BLT_2_R has been suggested to promote resolution of inflammation [9]. Furthermore, BLT_2_R is widely expressed in human tissues and is a low affinity BLTR that also binds other AA-derived molecules, whereas BLT_1_R is a high affinity BLTR that is predominantly expressed in human leukocytes, including neutrophils [11-14]. LTB_4_ is a well-characterised neutrophil chemoattractant that potently promotes migration [15], secretion of granule components (degranulation) and an increase in the intracellular concentration of free calcium ions ([Ca^2+^]_i_) through activation of BLT_1_R [9, 16]. Previous research has shown increased blood levels of LTB_4_ in certain inflammatory conditions such as rheumatoid arthritis and asthma [17, 18]. Increases in LTB_4_ can occur in response to a variety of stimuli [19, 20], which activate various enzymes (e.g., phospholipase A_2_ and 5-lipoxygenase) that mediate intracellular biosynthesis prior to transport (by an as yet unknown carrier protein) and release into the extracellular milieu ([9, 16, 17, 21]). Human neutrophils are a major source of LTB_4_, which can promote inflammation, for example through a paracrine mechanism leading to neutrophil swarming [20, 22, 23]. Many neutrophil chemoattractants that mediate migration, degranulation and elevated [Ca^2+^]_i_ also trigger the production of NADPH oxidase-derived reactive oxygen species (ROS) upon binding to their cognate GPCRs [7]. However, this is not the case for LTB_4_, which has been described as a biased/functionally selective neutrophil agonist with a weak capacity to activate the NADPH oxidase [24-29]. The ROS generated by the superoxide anion (O_2_^-^; the precursor of all ROS) producing NADPH oxidase system are critical to our host defence due to their ability to kill invading pathogens. However, because ROS are destructive/toxic not only to microbes but also to host cells and tissues, the NADPH oxidase activation must be properly regulated [30]. Under healthy conditions, this regulation is partly maintained by neutrophils circulating in the bloodstream in a quiescent (naïve) state, producing very low levels of GPCR-mediated NADPH oxidase-derived ROS. One reason for this is that most plasma membrane receptors (e.g., GPCRs including the formyl peptides [FPRs]) in naïve neutrophils are localised in the intracellular granules [31]. However, upon exposure to certain mediators (e.g., lipopolysaccharides [LPS] and/or tumour necrosis factor α [TNFα]), neutrophils can be induced to enter a pre-activated/primed state characterised by enhanced NADPH oxidase activity when triggered by certain GPCR agonists [7, 32, 33]. Primed neutrophils are often (but not always) characterised by increased receptor exposure on the plasma membrane (as a result of receptor mobilisation from granule stores), but some priming agents (e.g., hyaluronan) selectively mediate enhanced NADPH oxidase activity and not increased receptor exposure [34, 35]. We have also recently uncovered a novel priming mechanism in which the free fatty acid receptor 2 (FFA2R) plays a key role in enhancing NADPH oxidase activity. This receptor is a GPCR expressed on neutrophils for which short chain fatty acids (SCFAs) such as acetate and propionate are agonists. When neutrophils are stimulated with SCFAs, downstream signals include an increased [Ca^2+^]_i_ but no activation of the NADPH oxidase. However, the presence of a non-activating positive and selective allosteric FFA2R modulator (compound 58 [Cmp58]) alters this activation profile. Consistent with the function of an allosteric GPCR modulator, the response induced in neutrophils by SCFAs is enhanced in the presence of the allosteric FFA2R modulator. However, the allosteric FFA2R modulator also converts other GPCR agonists, unrelated to the FFA2R, into potent neutrophil activators. These data not only add this receptor transactivation process as a novel priming mechanism, but also break the receptor restriction properties that normally define the selectivity of allosteric GPCR modulators [30, 36-38].

It has not been investigated whether this novel neutrophil priming mechanism, based on a receptor transactivation between FFA2R and an unrelated GPCR, also occurs when the unrelated GPCR is BLT_1_R. Therefore, in this study we set out to investigate whether allosteric modulation of FFA2R could also convert LTB_4_ from a weak to a potent activator of the neutrophil NADPH oxidase. Our results not only confirm previous data showing that LTB_4_ alone is a weak activator of NADPH oxidase [24-29], but also show that LTB_4_ can indeed be converted to a potent activator of the NADPH oxidase in the presence of the allosteric FFA2R modulator Cmp58. Similar to previously described receptor transactivation between allosterically modulated FFA2Rs and other (non-FFA2R) agonist-bound GPCRs, the FFA2R-BLT_1_R transactivation was also functionally selective (biased) towards NADPH oxidase activation, as the LTB_4_-induced increase in [Ca^2+^]_i_ was unaffected by the presence of Cmp58. The data presented in this study therefore provide further evidence to re-evaluate the commonly accepted notion of allosteric modulation, which assumes that an allosteric receptor modulator only affects the response induced by orthosteric agonists that are specifically recognised by the receptor that binds the allosteric modulator.

## 2. Material and Methods

### 2.1. Chemicals and reagents

Dextran T500 (#551005009007) was purchased from Pharmacosmos (Holbaek, Denmark) and Cytiva Ficoll-Paque^TM^ Plus Media (#17-1440-03) was from Fischer Scientific (Gothenburg, Sweden). LTB_4_ (# 20110) and the LTB_4_ receptor antagonist U-75302 targeting BLT_1_R (# 70705) were from Cayman Chemical (Ann Arbor, MI, USA). Isoluminol (# A-8264), TNFα (# T6674), the FPR1 agonist fMLF (*N*-formyl-methionyl-leucyl-phenylalanine, #F3506), the FFA2R agonist propionic acid (propionate; # 94425) and the positive allosteric FFA2R modulator Cmp58 ((*S*)-2-(4-chlorophenyl)-3,3-dimethyl-*N*-(5-phenylthiazol-2-yl)butanamide; # 371725) were from Sigma-Aldrich (Merck, Burlington, MA, USA). Horseradish peroxidase (HRP, peroxidase [POD], #10108090001) and bovine serum albumin fraction V (BSA, #10735094001) were from Roche (Merck, Burlington, MA, USA) whereas Fura-2-acetoxymethyl ester (AM, #F1221) was from Invitrogen by Thermo Fischer Scientific (Gothenburg, Sweden). The Gα_q_ inhibitor YM-254890 (#257-00631) was purchased from FUJIFILM Wako Chemicals Europe (Nordic Biolabs, Täby, Sweden) and the FFA2R antagonist CATPB ((*S*)-3-(2-(3-Chlorophenyl)acetamido)-4-(4-(trifluoromethyl) phenyl) butanoic acid; # 5903) was from Tocris Bioscience (Bristol, UK).

All ligand stocks were dissolved, aliquoted and stored as recommended by the manufacturer and subsequent dilutions were made in Krebs-Ringer glucose (KRG) buffer (120 mM NaCl, 5 mM KCl, 1.5 mM MgSO_4_, 1.7 mM KH_2_PO_4_, 8.3 mM Na_2_HPO_4_, 10 mM glucose in the absence or presence of 1 mM CaCl_2_ [KRG^-^ and KRG^+^, respectively] in dH_2_O, pH 7.3) on the day of the experiment.

### 2.2. Ethical statement

This study involves human neutrophils isolated from buffy coats obtained from by healthy blood donors at the blood bank at Sahlgrenska University Hospital, Gothenburg, Sweden. As the buffy coats were donated anonymously and terefor could not be traced back to a specific donor, no ethical approval was required according to the Swedish legislation section code 4§ 3p SFS 2003:460 (Law on Ethical Testing of Research Relating to People).

### 2.3. Isolation of human neutrophils

Neutrophils were isolated from buffy coat blood obtained from healthy blood donors according to the method developed by Bøyum [39-41] and described in detail in our previous publication [41]. Briefly, neutrophils were isolated from the buffy coat blood by dextran sedimentation and Ficoll-Paque gradient centrifugation. Residual erythrocytes were removed by hypotonic lysis and washing in KRG before analysis of the neutrophil purity and concentration on a Sysmex KX-21N Haematology Analyzer (Sysmex Corporation, Kobe, Japan) or an Accuri C6 flow cytometer (Becton Dickinson, San Jose, CA, USA. Isolated neutrophils were diluted to a concentration of 1x10^7^ neutrophils/mL in KRG^+^ and stored on ice until used in experiments performed on the day of isolation. This study includes 32 different blood donors and the purity (mean ± SD) of neutrophils isolated from the buffy coats of these individuals was 94 % ± 2 %.

### 2.4. Measurement of alterations in the intracellular concentration of free calcium ions

Fura 2-AM loaded neutrophils were analysed for an agonist mediated rise in intracellular calcium ion concentration [Ca^2+^]_i_ by fluorescence spectrophotometry as described in detail in our previous publication [41]. In brief, isolated neutrophils (2 × 10^7^ cells/ml in KRG^-^ containing 0.1% BSA) were stained with Fura 2-AM (2 µg/mL, 30 min at room temperature in darkness) and thereafter washed twice before resuspended to concentration of 2x10^7^ neutrophils/mL in ice cold KRG^+^. The Fura 2-AM stained neutrophils, were kept on ice in darkness until measurements of [Ca^2+^]_i_ on a Perkin Elmer fluorescence spectrophotometer (LC50B), with excitation wavelengths of 340 nm and 380 nm, an emission wavelength of 509 nm, and slit widths of 5 nm and 10 nm, respectively. For the measurements, disposable polystyrene cuvettes (4.2 mL) containing 2.475 mL reaction mixture (KRG^+^ and 5 x 10^6^ Fura 2-AM stained neutrophils) were first equilibrated at 37°C for ten minutes, prior addition of an agonist (25 µL). For analysis of how the presence of U-75302, YM-254890, CATPB or Cmp58 affected the agonist mediated rise in [Ca^2+^]_i_, these compounds were added to the reaction mixture just before the equilibration at 37°C for ten minutes. The transient increase in [Ca^2+^]_i_ data is shown as the ratio of detected fluorescence intensities (340 nm: 380 nm) detected. For summary analyses, the peak [Ca^2+^]_i_ was used. This was calculated by subtracting the background level, defined as the peak value measured just prior to stimulation with the respective agonist from the measured agonist-induced peak [Ca^2+^]_i._

### 2.5. Measurement of NADPH oxidase derived-superoxide anions

To measure the release of superoxide anions (O_2_; the precursors of reactive oxygen species [ROS]), generated upon activation of the neutrophil NADPH oxidase, an isoluminol-amplified chemiluminescence (CL) system was utilized as previously described [42, 43]. Briefly, disposable polypropylene tubes (4 mL) containing 0.9 mL of a reaction mixture (KRG^+^, 4 U/mL HRP, 10 µg/mL isoluminol and 10^5^ isolated neutrophils) were initially equilibrated at 37°C for five minutes, prior addition of an agonist (100 µL) and measurement of O_2_^-^ production/light emission over time in a six-channel Biolumat LB 9505 (Berthold Co., Wildbad, Germany). For some experiments the isolated neutrophils (1 x 10^6^/mL) were treated (primed) with TNFα (10 ng/mL, 37°C, 20 minutes) and then stored on ice prior added to the reaction mixture for measurement of O_2_ production. For analysis of how the presence of U-75302, YM-254890, CATPB or Cmp58 affected the agonist-induced O_2_ production, these compounds were added to the reaction mixture just before the equilibration at 37°C for five minutes. The O_2_ production/light emission is expressed as is expressed as Mega counts per minute (Mcpm). For summary analyses of O_2_ production the area under the curve (AUC) was used. This was calculated by subtracting the background level, defined as the AUC measured prior to stimulation with the respective agonist, from the measured agonist-induced AUC of the O_2_ production.

## Data analysis

Data analysis was conducted using GraphPad Prism 10 for macOS, version 10.2.3 (GraphPad Software, San Diego, CA, USA). Each experiment was conducted a minimum of three times with isolated neutrophils from different healthy blood donors. For data were the half maximal effective concentration (EC_50_) of a ligand was determined the curve fitting was achieved by non-linear regression using the sigmoidal dose–response equation (variable-slope). All statistical calculations were performed using raw data values using a paired two-tailed Student’s *t*-test for data comparing two groups and an ordinary or repeated measures one-way ANOVA followed by either Sidak’s or Dunnett’s multiple comparisons test for data comparing more than two groups. Where applicable, the specific statistical tests used for each figure are described in the figure legends. Statistically significant differences are indicated by \**p* < 0.05, \*\**p* < 0.01, \*\*\**p* < 0.001 and \*\*\*\**p* < 0.0001, no statistically significant differences (*p* > 0.05) are not indicated.

## 3. Results

### 3.1. LTB_4_ is a potent inducer of the transient rise in cytosolic concentration of free calcium ions but a weak activator of the NADPH oxidase

The two neutrophil chemoattractants LTB_4_ and the *N*-formylated tripeptide fMLF [44], induce a phospholipase C (PLC) mediated increase of the intracellular concentration of free calcium ions ([Ca^2+^]_i_) in neutrophils (Fig 1A and [45]). This PLC – phosphatidyl inositol-4,5-*bis*-phosphate (PIP_2_) – inositol-1,4,5-*tris*-phosphate (IP_3_) mediated rise in [Ca^2+^]_i_ is one of the earliest downstream signalling events following an agonist activation of neutrophil GPCRs [36]. When comparing the LTB_4_ and fMLF mediated increase in [Ca^2+^]_i_, our results revealed a similar increase in [Ca^2+^]_i_ when triggering neutrophils with 10 nM of either of these two chemoattractants (Fig. 1A). When using concentrations < 10 nM, our data show that LTB_4_ is a slightly more potent inducer of [Ca^2+^]_i_ than fMLF (Fig. 1A-C). Furthermore, the LTB_4_ response was completely inhibited by the BLT_1_R antagonist U-75302 (Fig. 1B-C), demonstrating that the LTB_4_ mediated transient rise in [Ca^2+^]_i_ is achieved through this high affinity LTB_4_ receptor [11, 46]. Also, and in agreement with the well-established receptor preference for fMLF, the fMLF mediated rise in [Ca^2+^]_i_ was fully inhibited by the recently described powerful FPR1 antagonist AZ2158 [47]. Apart from mediating a transient rise in [Ca^2+^]_i_, fMLF is also a potent activator of the superoxide anion (O_2_^-^) generating NADPH oxidase [7]. This contrasts with LTB_4_, which capacity to activate the NADPH oxidase has been demonstrated to be rather weak as compared to other neutrophil chemoattractants, including fMLF [24-29]. Our data confirm these previous finding and show that while fMLF is a potent inducer of ROS, LTB_4_ is a very weak inducer of this neutrophil effector function (Fig. 2A – B). This weak capacity of LTB_4_ to mediate production of ROS was regardless of the neutrophil state (naïve or primed with the classical priming agent TNFα), whereas the fMLF response, run in parallel, was largely increased in TNFα-primed as compared to naïve neutrophils (Fig. 2A – B). That is, despite LTB_4_ being slightly more potent than fMLF at mediating a rise in [Ca^2+^]_i_, an effect achieved through activation of BLT_1_R, this lipid is a biased signalling agonist with a weak capacity to activate the NADPH oxidase.

**Figure 1.**
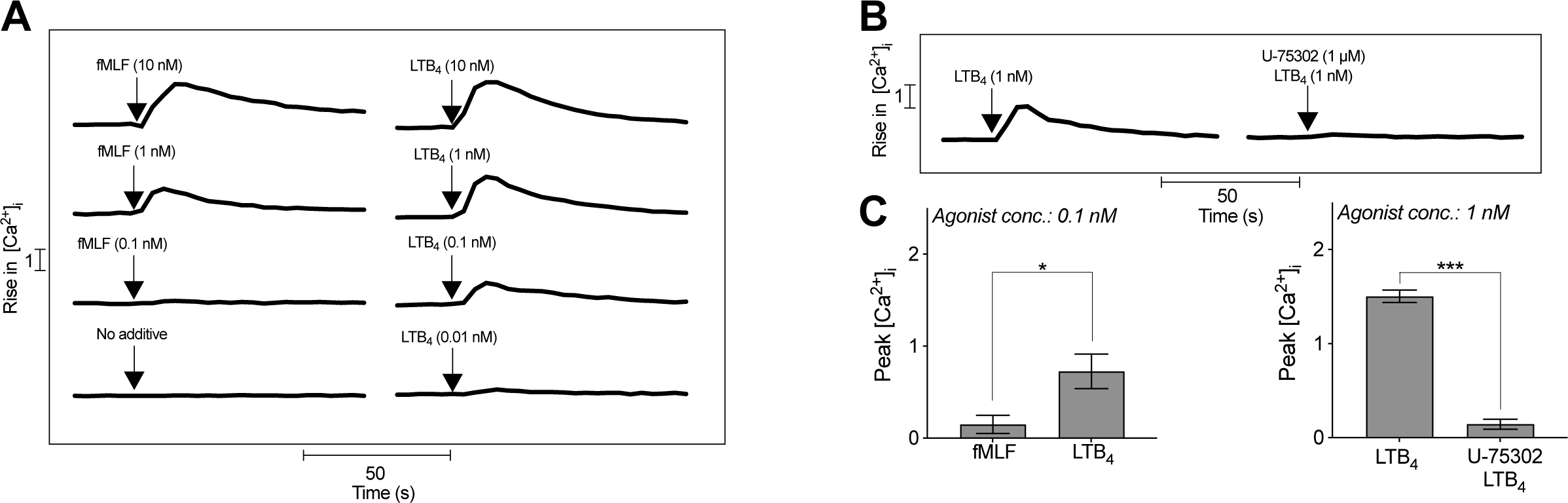
LTB_4_ induces a transient rise in intracellular Ca^2+^ in neutrophils that rely on activation of BLT_1_R. Fura 2-AM loaded neutrophils were pre-incubated for ten min at 37°C in the absence or presence of the BLT_1_R antagonist U-75302 prior to stimulation with fMLF or LTB_4_ and measurement of a rise in the free intracellular calcium ion (Ca^2+^) concentration [Ca^2+^]_i_ over time. **A**) Neutrophils were stimulated (time of agonist addition is indicated by an arrow) with different concentrations of fMLF (10, 1 or 0.1 nM; left panel), buffer (no additive; left panel) or LTB_4_ (10, 1, 0.1, 0.01 nM; right panel). One representative out of 3-4 individual experiments for each agonist concentration is shown. **B**) Neutrophils were pre-incubated in the absence or presence of U-75302 (1 µM) prior stimulation (time of agonist addition is indicated by an arrow) with LTB_4_ (1 nM). One representative out of 3 individual experiments for each agonist concentration is shown. **C**) Comparison of the peak [Ca^2+^]_i_ induced by 0.1 nM fMLF or 0.1 nM LTB_4_ (mean ± SEM, n = 4, left bar graph) or by 1 nM LTB_4_ in the absence or presence of 1 µM U-75302 (mean ± SEM, n = 3, right bar graph). Statistically significant differences for the data in each bar graph were evaluated by a paired Student’s *t*-test (\**p*<0.05, \*\*\**p*<0.001).

**Figure 2.**
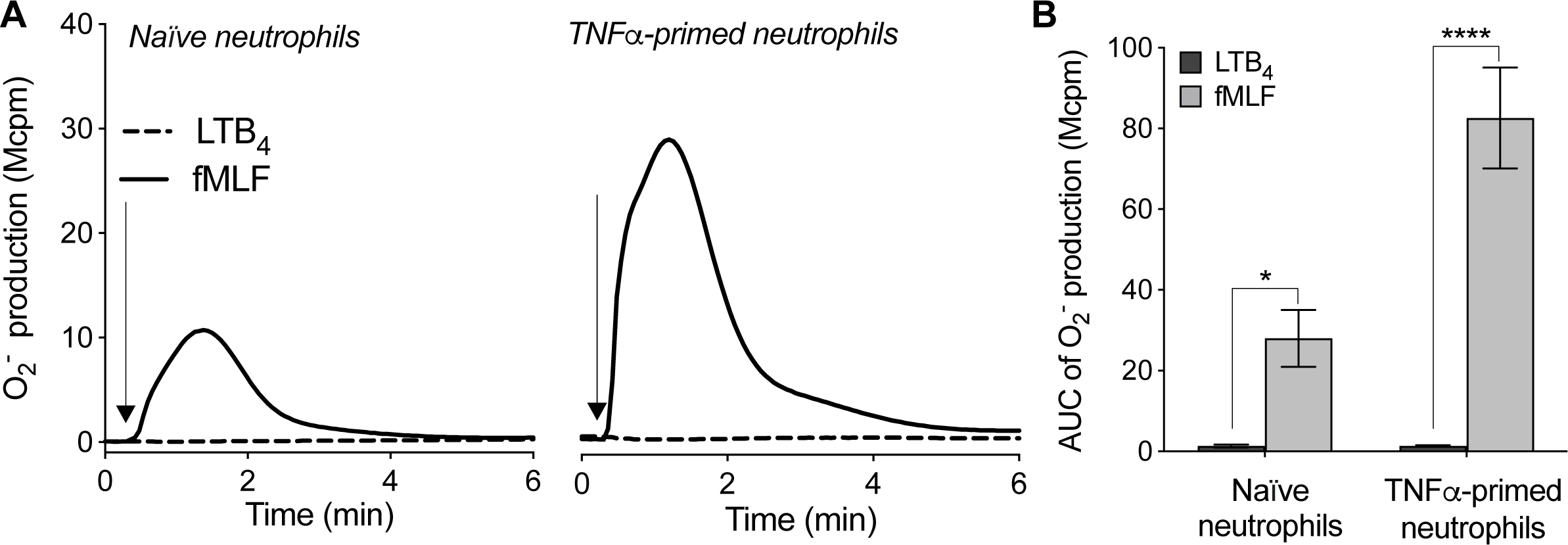
LTB_4_ is a weak inducer of NADPH oxidase activity in both naïve and TNFα-primed neutrophils. Neutrophils were incubated without (naïve) or with TNFα (10 ng/mL, 37°C, 20 minutes; TNFα-primed) prior stimulation with LTB_4_ (10 nM) or fMLF (10 nM) and measurement of NADPH oxidase produced oxygen radicals (O_2_^-^) over time. **A**) Representative traces of naïve (left panel) and TNFα-primed (right panel) neutrophils stimulated (time of agonist addition is indicated by an arrow) with either LTB_4_ or fMLF. **B**) The bar graph shows the calculated area under the curve (AUC) of the O_2_^-^ produced by the two agonists in naïve and TNFα-primed neutrophils (mean ± SEM, n = 5). Statistically significant differences were evaluated by repeated measures one-way ANOVA with Sidak’s multiple comparisons test for analysis of the fMLF and LTB4 induced responses in naïve and TNFα-primed neutrophils, separately (\**p* < 0.05, \*\*\*\**p* < 0.0001).

### 3.2. LTB_4_ is converted into a potent activator of the neutrophil NADPH oxidase in the presence of the allosteric FFA2R modulator Cmp58

Propionate, a short chain fatty acid produced by the gut microbiota during fermentation of dietary carbohydrates, is an orthosteric agonist recognized be FFA2R. We have previously shown, that in human neutrophils, propionate acts as a biased signalling agonist, that like LTB_4_ induces a transient rise in [Ca^2+^]_i_, whereas it’s capacity to activate the NADPH oxidase is rather scarce. However, presence of the positive allosteric FFA2R modulator termed Cmp58, which by itself does not have any direct neutrophil activating effect, turns propionate into a potent NADPH oxidase activator in TNFα primed neutrophils (Fig 3A-B and [38, 48]). These data thus agrees with the dogma for how the response induced by an orthosteric agonist should be increased in the presence of a positive allosteric modulator recognized by the same receptor but having a binding site spatially different from the orthosteric site [36]. However, presence of Cmp58 also turned LTB_4_ into a potent NADPH oxidase activating agonist with a time-course and magnitude of the LTB_4_ mediated response quite similar to that induced by propionate in the presence of Cmp58 (Fig 3A-B). This effect of Cmp58 was most prominent in TNFα primed neutrophils in which LTB_4_ displayed a half maximal effective concentration (EC_50_) of around 9 nM (Fig 3C). It should however also be noted that presence of Cmp58 also positively affected LTB_4_’s capacity to induce NADPH oxidase derived ROS in naïve neutrophils, although the magnitude of the response was lower as compared to that induced in TNFα primed neutrophils presence of Cmp58 (Fig. 3A; inset). This is in line with our previous data showing that Cmp58’s capacity to turn orthosteric FFA2R agonists and the P2Y_2_R agonist ATP into potent NADPH oxidase activators is most pronounced in TNFα primed as compared to naïve neutrophils [38, 48]. Taken together, these data show that the lipid agonist LTB_4_ is turned from a weak to a strong activator of the neutrophil NADPH oxidase provided that FFA2R has been allosterically modulated prior to stimulation with LTB_4_.

**Figure 3.**
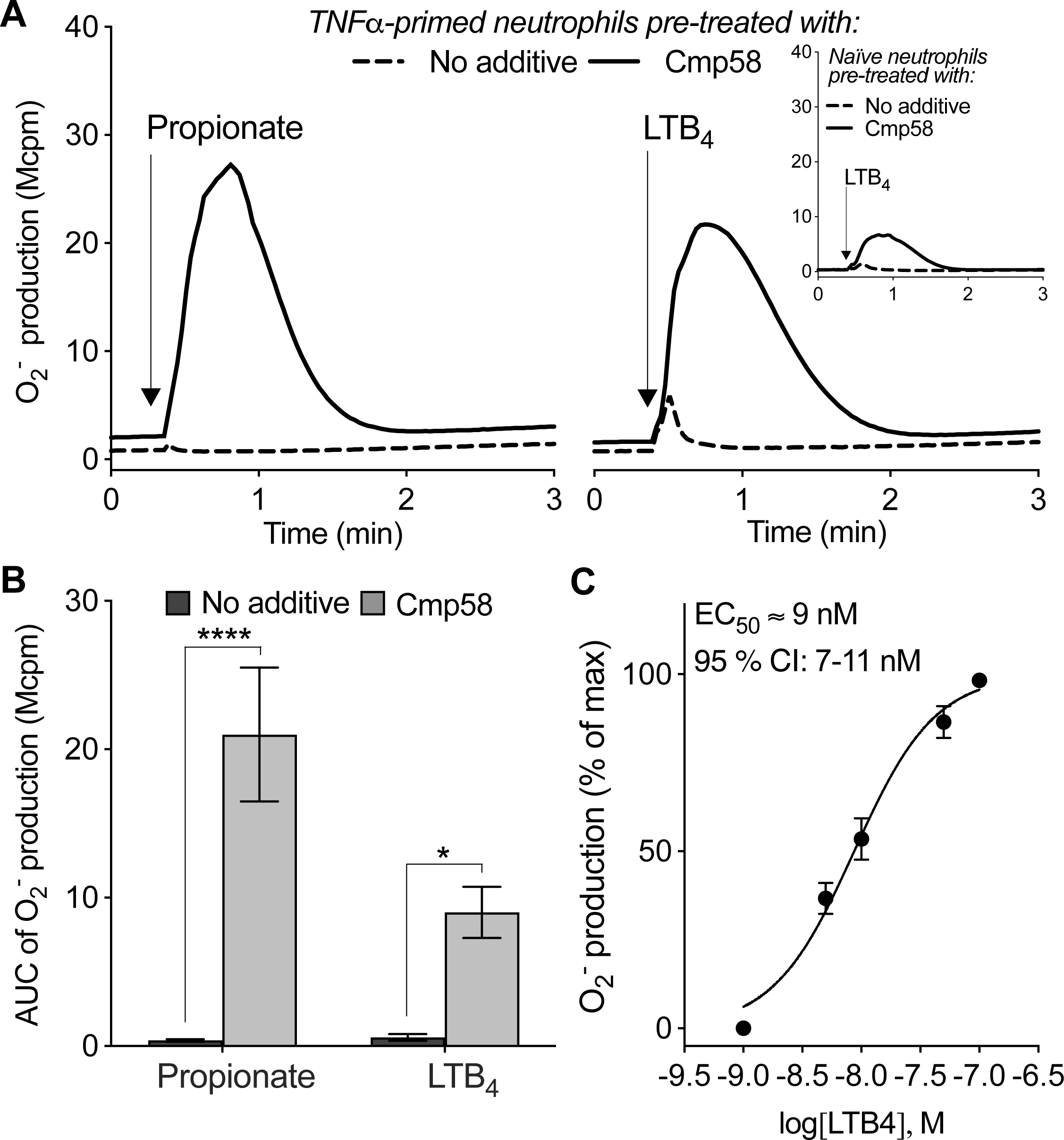
LTB_4_ is turned into a potent NADPH oxidase activator in TNFα-primed neutrophils pre-treated with the allosteric FFA2R modulator Cmp58. Neutrophils were incubated without (naïve) or with TNFα (10 ng/mL, 37°C, 20 minutes; TNFα-primed) and then pre-treated in the absence (no additive) or presence of the FFA2R allosteric modulator Cmp58 (1 µM) at 37°C for five minutes prior stimulation with an agonist and measurement of NADPH oxidase produced oxygen radicals (O_2_^-^) over time. **A**-**B**) Neutrophils were stimulated with the FFA2R orthosteric agonist propionate (25 µM) or LTB_4_ (10 nM). **A**) Representative traces of TNFα-primed neutrophils stimulated with propionate (left panel) or LTB_4_ (right panel; time of agonist addition is indicated by an arrow). **Inset**: Representative traces of naïve neutrophils stimulated with LTB_4_ (time of agonist addition is indicated by an arrow; one representative out of 4 individual experiments for each agonist concentration is shown). **B**) The bar graph shows the calculated area under the curve (AUC) of the O_2_ ^-^ produced by the two agonists in TNFα-primed neutrophils pre-treated in the absence or presence of Cmp58 (mean ± SEM, n = 7). Statistically significant differences in B were evaluated by a repeated measures one-way ANOVA with Sidak’s multiple comparisons test for analysis of the propionate and LTB4 induced responses in TNFα-primed neutrophils pre-treated with or without Cmp58, separately (\*\*\*\**p* < 0.0001). **C**) TNFα-primed neutrophils pre-treated with Cmp58 were stimulated with different concentrations of LTB_4_. A dose-response curve of the LTB_4_-induced O_2_ production, including the half maximal effective concentration (EC_50_) of LTB_4_ and the 95 % confidence interval (CI) of the EC_50_ value (mean ± SEM, n = 6).

### 3.3. The LTB_4_ induced ROS production relies on a receptor transactivation mechanism involving FFA2R – the response is inhibited by presence of a FFA2R antagonist or a BLT_1_R antagonist but unaffected by an inhibitor of Gα_q_

The LTB_4_ mediated ROS production was initiated very rapidly after addition of the agonist, and the peak of the response was reached within a minute after addition (Fig 3A). As the LTB_4_ mediated ROS production was dependent not only on LTB_4_ but also on the presence of the allosteric FFA2R modulator Cmp58 (Fig 3A-B), our data strongly suggest that the response is mediated by a receptor transactivation mechanism between the two receptors BLT_1_R and FFA2R. To investigate such a receptor transactivation in more detail we next tested the sensitivity of the LTB_4_ induced response to receptor selective antagonists. Our results show that not only the BLT_1_R antagonist U-75302 potently reduced the LTB_4_ mediated NADPH oxidase activity, but also the FFA2R specific antagonist CATPB inhibited the response (Fig 4A-B). This strongly support the suggestion that a novel receptor transactivation mechanism by which signals generated by the LTB_4_ activated BLT_1_Rs activate the allosterically modulated FFA2Rs, a mechanism that allows for LTB_4_ to potently activate the ROS producing NADPH oxidase in neutrophils. Our previous work has shown that the Ga_q_ inhibitor YM-254890 inhibits activation of the human neutrophil NADPH oxidase when activated by some GPCR agonists e.g., platelet activating factor (PAF) binding to the PAF receptor, but not others e.g., fMLF which bind to FPR1 (Fig 5A-B and [7, 49]). As our results are the first to demonstrate LTB_4_ as a potent NADPH oxidase activator provided that FFA2R has been allosterically modulated prior to stimulation, we next investigated if this response was sensitive to Ga_q_ inhibition. However, the Ga_q_ inhibitor YM-254890 did not affect the response (Fig 5C-D) and we can therefore conclude that this GPCR transactivation between BLT_1_R and FFA2R is not dependent on Ga_q_. Taken together, these data show that the ROS production mediated by LTB_4_ in TNFα primed neutrophils with allosterically modulated FFA2Rs can be inhibited by both a selective BLT_1_R antagonist and a selective FFA2R antagonist, but not by the Ga_q_ inhibitor YM-254890. Hence, these results demonstrate that a novel GPCR transactivation mechanism between BLT_1_R and FFA2R turns LTB_4_ into a potent neutrophil NADPH oxidase activating agonist.

**Figure 4.**
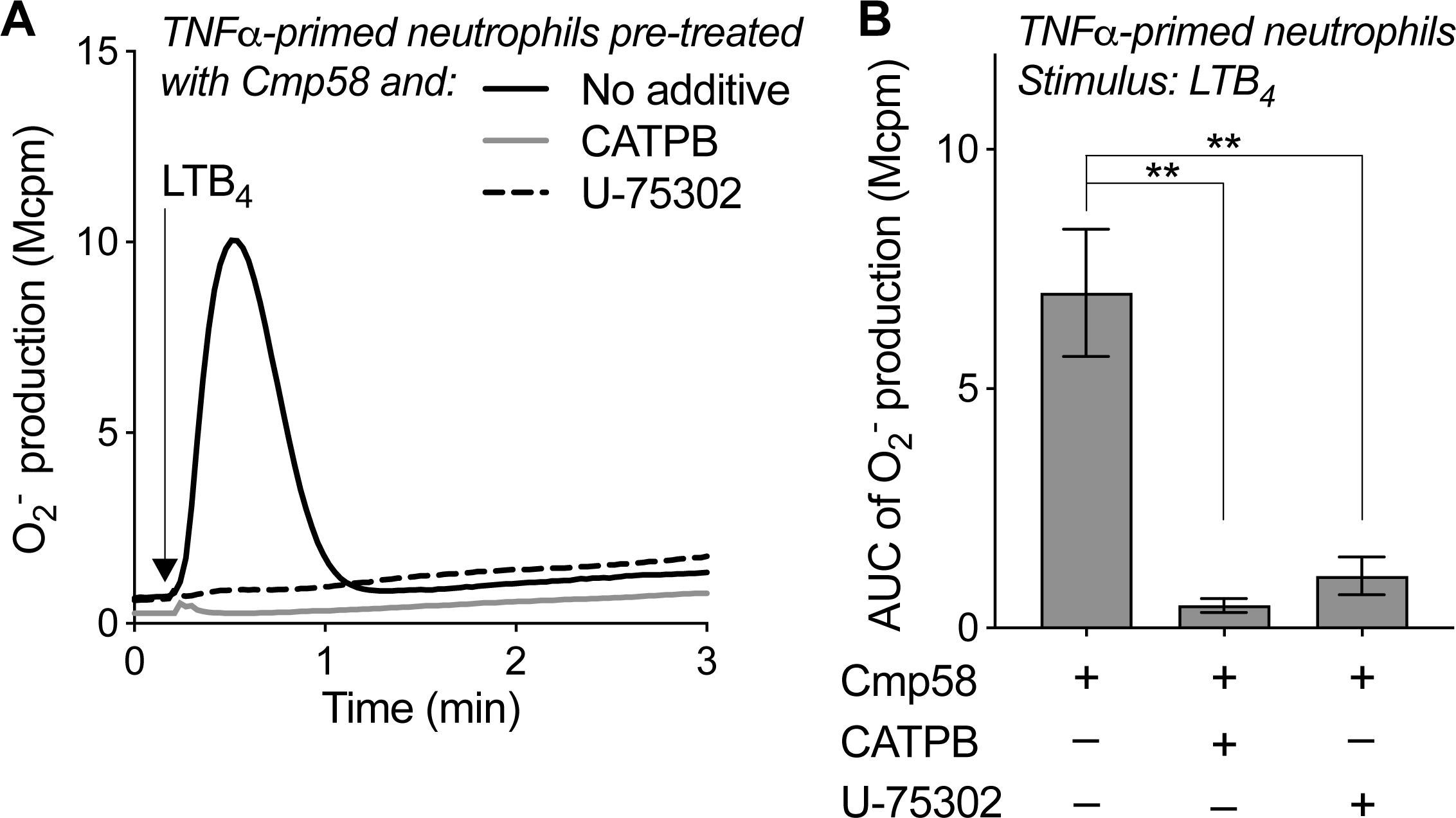
The LTB_4_ mediated NADPH oxidase activity by TNFα-primed neutrophils pre-treated with Cmp58 can be inhibited by either BLT_1_R or FFA2R selective antagonists. Neutrophils were incubated with TNFα (10 ng/mL, 37°C, 20 minutes; TNFα-primed) and then pre-treated with the FFA2R allosteric modulator Cmp58 (1 µM) in the absence (no additive) or presence of the BLT_1_R receptor antagonist U-75302 (1 µM) or the FFA2R antagonist CATPB (0.1 µM) for five minutes at 37°C. Thereafter the neutrophils were stimulated with LTB_4_ (10 nM), and NADPH oxidase produced oxygen radicals (O_2_^-^) was measured of over time. **A**) Representative traces of TNFα-primed neutrophils pre-treated with Cmp58 in the absence or presence of U-75302 or CATPB prior stimulation with LTB_4_ (time of stimulation is indicated by an arrow). **B**) The bar graph shows the calculated area under the curve (AUC) of the O_2_ produced by LTB_4_ in TNFα-primed neutrophils pre-treated with Cmp58 in the absence or presence of U-75302 or CATPB prior stimulation with LTB_4_ (mean ± SEM, n = 6). Statistically significant differences were evaluated by repeated measures one-way ANOVA and Dunnett’s multiple comparisons test to the no additive control (\*\**p* < 0.01).

**Figure 5.**
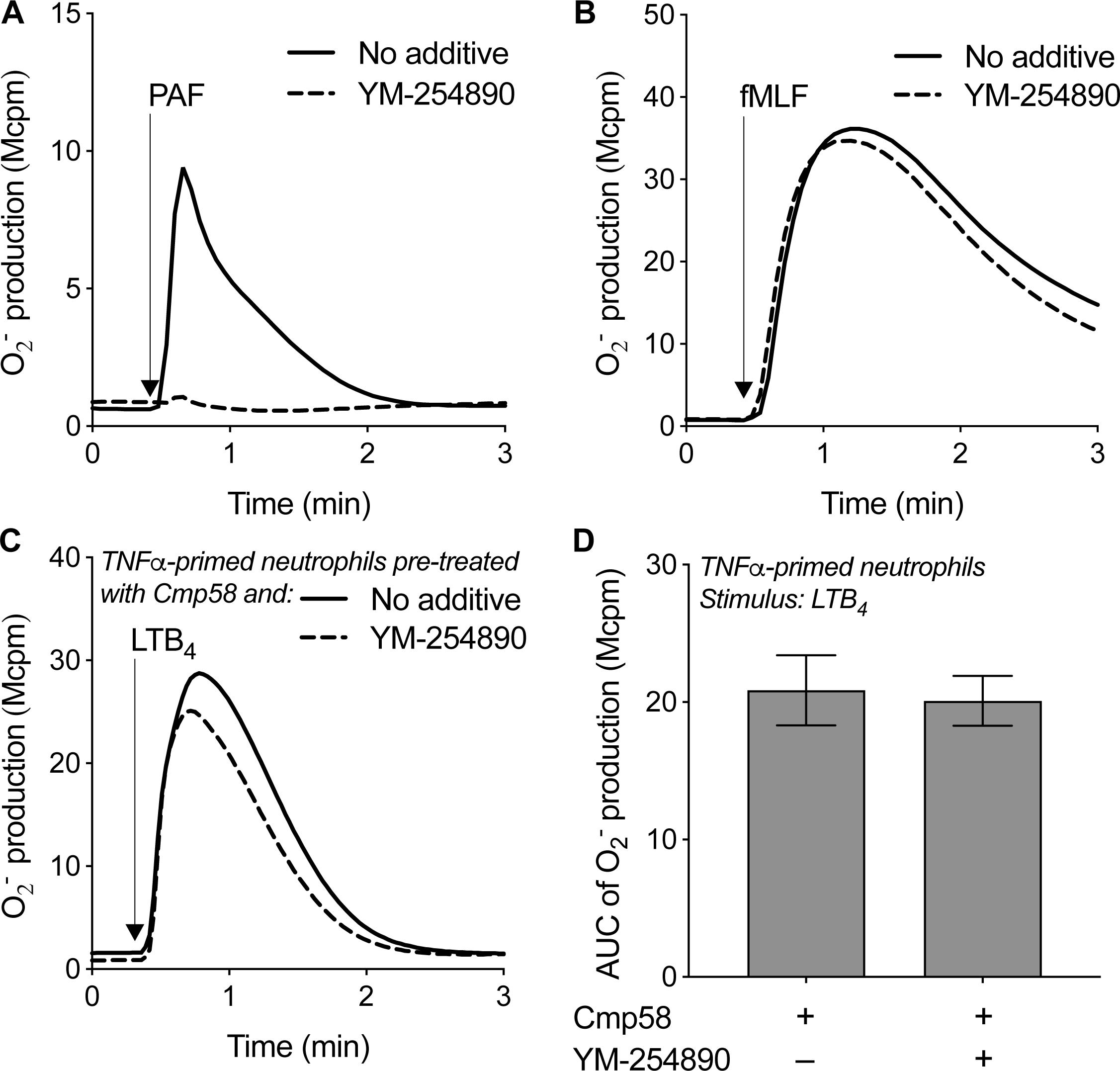
The LTB_4_ mediated NADPH oxidase activity by TNFα-primed neutrophils pre-treated with Cmp58 does not engage Gαq proteins for signal transduction. Neutrophils were pre-treated in the absence (no additive) or presence of the Gαq inhibitor YM-254890 (200 nM) for five minutes at 37°C prior agonist stimulation and measurement of NADPH oxidase produced oxygen radicals (O_2_^-^) over time. **A**-**B**) Representative traces of naïve neutrophils pre-treated with or without YM-254890 for five minutes at 37°C prior stimulation (time of agonist addition is indicated by an arrow) with **A**) platelet activating factor (PAF, 100 nM) or **B**) fMLF (100 nM). One representative out of three individual experiments for each agonist is shown. **C**-**D**) Neutrophils were incubated with TNFα (10 ng/mL, 37°C, 20 minutes; TNFα-primed) before pre-treated for five minutes at 37°C with the FFA2R allosteric modulator Cmp58 (1 µM) in the absence or presence of YM-254890 prior stimulation with LTB_4_ (10 nM). **C**) Representative traces of the LTB_4_-induced O_2_ production (time of LTB_4_ stimulation is indicated by an arrow). **D**) The bar graph shows the calculated area under the curve (AUC) of the O_2_ produced by LTB_4_ in TNFα-primed neutrophils pre-treated with Cmp58 in the absence or presence of YM-254890 prior stimulation (mean ± SEM, n = 5). Statistically significant differences were evaluated by a paired Student’s *t*-test (*p* = 0.76).

### 3.4. The receptor transactivation mechanism between BLT_1_R and FFA2R, that mediates LTB_4_-induced ROS production, is functionally selective

As LTB_4_ was such a weak inducer of NADPH oxidase derived ROS in the absence of Cmp58 (Fig 2 and 3) we utilized LTB_4_’s capacity to mediate a transient rise in [Ca^2+^]_i_ in naïve neutrophils (Fig 1) to further evaluate the GPCR transactivation between BLT_1_R and FFA2R. We have previously shown that the orthosteric FFA2R agonist propionate can mediate a transient rise in [Ca^2+^]_i_ in naïve neutrophils and that this response is augmented in the presence of Cmp58, an effect that was especially clear when using low (suboptimal) propionate concentrations that barely mediate any increase in [Ca^2+^]_i_ in the absence of Cmp58 [38]. However, Cmp58 had no effect on the LTB_4_ mediated increase in [Ca^2+^]_i_; this was evident when using two suboptimal LTB_4_ concentrations (1 nM and 0.1 nM, Fig 1 and Fig 6A-B). In accordance with the lack of effect of Cmp58 on the LTB_4_ mediated increase in [Ca^2+^]_i_, also presence of the FFA2R antagonist CATPB lacked effect on the LTB_4_ mediated increase in in [Ca^2+^]_i_ (Fig 6A-B). The lack of an amplifying effect on the LTB_4_ mediated increase in [Ca^2+^]_i_ of the allosteric FFA2R modulator Cmp58, was thus in stark contrast to the Cmp58 dependent augmentation of the NADPH oxidase activity; in accordance with the Cmp58 dependency of the significantly augmented NADPH oxidase activity, this response was inhibited by CATPB (Fig. 3-4). Based on the lack of effect of the FFA2R ligands on the LTB_4_ mediated increase in [Ca^2+^]_i_ we conclude that the signals generated by the BLT_1_R induced transactivation of FFA2R that leads to NADPH oxidase activity are functionally selective and only positively affects the pathway that leads to an activation of the NADPH oxidase while leaving the PLC-PIP_2_-IP_3_-Ca^2+^ route unaffected. Moreover, the LTB_4_ mediated increase in [Ca^2+^]_i_ in the presence of Cmp58 was, similar to the NADPH oxidase activity (Fig 4-5), unaffected by presence of the Ga_q_ inhibitor YM-254890 but inhibited by the BLT_1_R antagonist U-75302 (Fig 6A-B). In summary, these results show that signalling following the BLT_1_R induced transactivation of the allosterically modulated FFA2R, that turns LTB_4_ into a potent NADPH oxidase activator is biased and selectively affect LTB_4_ mediated ROS production while leaving the PLC-PIP_2_-IP_3_-Ca^2+^ route leading to an increase in [Ca^2+^]_i_ unaffected.

**Figure 6.**
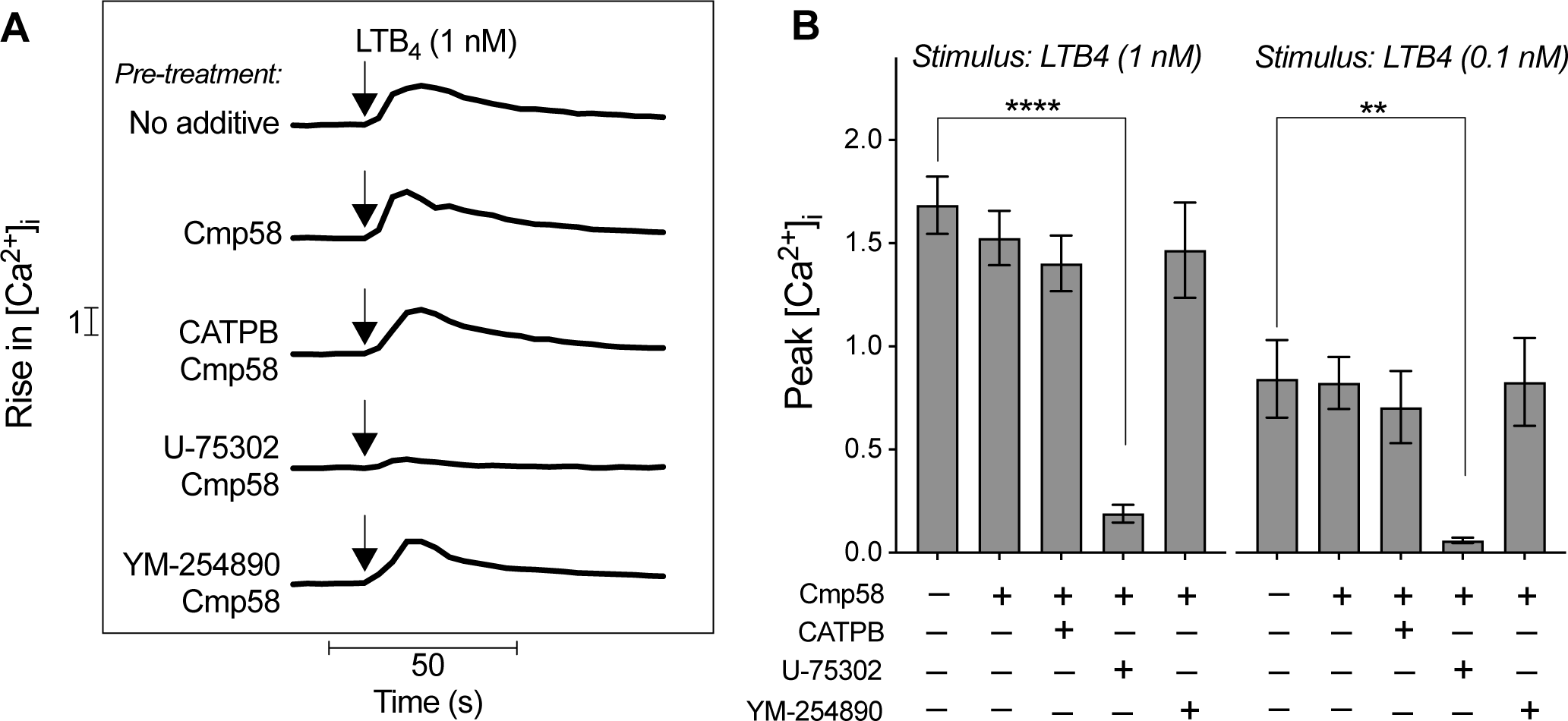
The LTB_4_-induced rise in intracellular calcium ion concentration is not affected by the allosteric FFA2R modulator Cmp58 and cannot be inhibited by CATPB. Fura 2-AM loaded neutrophils were pre-treated for ten min at 37°C in the absence (no additive) or presence of FFA2R allosteric modulator Cmp58 (1 µM), the FFA2R antagonist CATPB (0.1 µM), the LTB_4_ receptor antagonist U-75302 (1 µM) and/or the Gαq inhibitor YM-254890 (200 nM) prior stimulation with LTB_4_ and measurement of the rise in intracellular calcium ion (Ca^2+^) concentration [Ca^2+^]_i_ over time. **A**) Differently pre-treated neutrophils were stimulated (time of agonist addition is indicated by an arrow) with LTB_4_ (1 nM). One representative out of 3-4 individual experiments for each condition is shown. **B**) The bar graphs show the peak [Ca^2+^]_i_ measured in the differently pre-treated neutrophils stimulated with LTB_4_ (1 nM, left panel; 0.1 nM, right panel; mean ± SEM, n = 3-4 for both panels). Statistically significant differences were evaluated by an ordinary one-way ANOVA and Dunnett’s multiple comparisons test to the no additive control (\*\**p* < 0.01, \*\*\*\**p* < 0.0001).

## 4. Discussion

The endogenous lipid inflammatory mediator LTB_4_, generated from AA through the 5-lipoxygenase pathway [9, 16, 17, 21], has been demonstrated to be a potent activator of human neutrophils. The precise signalling pathways that are triggered and regulate the biological activities mediated upon LTB_4_ binding to BLT_1_R (the high-affinity GPCR for LTB_4_) remain unknown. It is evident, however, that LTB_4_ is a biased or functional selective agonist in naïve human neutrophils. This is illustrated by the observation that LTB_4_ is a highly potent neutrophil chemoattractant and inducer of the PLC-dependent elevation in [Ca^2+^]_i_ [9, 15, 16], whereas it is a relatively weak activator of the superoxide anion-generating NADPH oxidase (this study and [24-29]). The present study demonstrates that a positive allosteric modulator (Cmp58) selective for FFA2R not only alters the response elicited by the orthosteric FFA2R agonist propionate, but also enhances the NADPH oxidase activity induced by LTB_4_. This FFA2R-dependent response was generated by a receptor transactivation mechanism initiated by signals that are mediated downstream of the LTB_4_-occupied BLT_1_Rs. In accordance with our receptor transactivation model proposed, the BLT_1_R induced intracellular signals activate the allosterically modulated FFA2Rs from the cytoplasmic side of the plasma membrane. Subsequently, this receptor transactivation mediates the assembly and activation of the neutrophil NADPH oxidase. The data demonstrating that the LTB_4_-mediated NADPH oxidase activity is completely inhibited by both a BLT_1_R-specific antagonist and an FFA2R-specific antagonist provide the foundation for this receptor transactivation model.

The receptor transactivation with LTB_4_ as the activating agonist is consistent with previous findings indicating that other low- and non-activating receptor agonists may be transformed into potent neutrophil NADPH oxidase activating agonists. In other words, these agonists also function as potent activators of the NADPH oxidase when they bind to their cognate receptors on neutrophils with allosterically modulated FFA2Rs. The aforementioned agonists include ATP, PAF, and non-activating concentrations of formylated peptides [30, 36-38]. Altogether, these findings demonstrate that intracellular signals that transactivate allosterically modulated FFA2Rs can be generated downstream of several agonist-activated neutrophil-expressed GPCRs. In addition to the anticipated augmentation of neutrophil NADPH oxidase-derived ROS production induced by orthosteric FFA2R agonists in the presence of Cmp58 (as demonstrated in this study and previous research [37, 38, 48, 50]), Cmp58 also enhances the neutrophil NADPH oxidase activity mediated by several other GPCR agonists. In light of these findings, the prevailing view on of allosteric modulation, namely that both positive (potentiating) and a negative (inhibiting) allosteric receptor modulators should influence only the response elicited by orthosteric agonists that are specifically recognised by the receptor that binds the allosteric modulator” [36, 51-53], requires reassessment. In the meantime, the development of specific positive allosteric modulators for neutrophil-expressed GPCRs other than FFA2R is eagerly awaited. This would permit studies to ascertain whether this unconventional “priming feature” by receptor transactivation involving allosterically modulated FFA2Rs is specific to FFA2R or also occurs when orthosteric agonists are used to stimulate neutrophils with allosterically modulated GPCRs other than FFA2R. Moreover, it remains to be seen whether neutrophil-expressed GPCRs can transactivate membrane receptors belonging to another receptor family. Furthermore, it is yet to be determined whether neutrophil-expressed GPCRs possess the capacity to transactivate membrane receptors belonging to a different receptor family. Prior research has for example indicated that the activation of the purinergic receptor P2Y_2_R (a GPCR that binds ATP) mediates transactivation of epithelial growth factor receptors in both brain and cancer [54-56].

A common mechanism has been identified in all FFA2R-unrelated GPCR agonists that have been demonstrated to enhance neutrophil NADPH oxidase activity through receptor transactivation of allosterically modulated FFA2Rs. This mechanism involves a PLC-dependent increase in [Ca^2+^]_i_, which is a very early event following GPCR agonist binding, irrespective of whether the receptor couple to a G protein containing a Gα_i_ or a Gα_q_ subunit [7]. In line with the inhibitory effect of the Gα_i_ selective inhibitor pertussis toxin, which indicates that both BLT_1_R and FFA2R couple to Gα_i_-containing G proteins [9, 30, 41, 57], the LTB_4_-induced transactivation was found to be insensitive to the Gα_q_ selective inhibitor YM-254890. In contrast, P2Y_2_R couple to Gα_q_-containing G proteins, and the ATP-induced transactivation has been demonstrated to be inhibited by YM-254890 [7, 41]. The differing coupling of the G protein observed between the neutrophil GPCRs that have been identified to transactivate allosterically modulated FFA2Rs indicates that the transactivation signals are generated downstream of the G protein. Although Cmp58 alone does not induce an elevation in [Ca^2+^]_i_ [48], the fact that the signalling cascade mediated by all neutrophil-expressed GPCRs that transactivate allosterically modulated FFA2Rs includes an elevation in [Ca^2+^]_i_, suggests that this PLC-PIP_2_-IP_3_ dependent pathway may be part of the GPCR transactivation mechanism. One potential mechanistic model for how such a receptor transactivation might occur, is that the allosteric FFA2R modulator transfers the FFA2Rs to a state that is activated from within the plasma membrane, by a rise in [Ca^2+^]_i_. Given that the PLC–PIP_2_-IP_3_–Ca^2+^ signalling route downstream of the transactivating GPCRs is not directly affected by Cmp58 (as demonstrated in this and previous studies [7, 38]), it can be postulated that the allosteric FFA2R modulator transfers FFA2R to a state that is directly activated by a rise in [Ca^2+^]_i_. This mechanistic model is corroborated by the results of our recent study, which demonstrated that activation of the NADPH oxidase can be induced by either ionomycin (a Ca^2+^-selective ionophore) or thapsigargin (a drug that increases [Ca^2+^]_i_ through an inhibition of the Ca^2+^ATPase in the endoplasmic reticulum) provided that the FFA2Rs were allosterically modulated when the rise in [Ca^2+^]_i_ was induced by these tool compounds [50].

The enhanced prominence of the GPCR transactivation mechanism in TNFα-primed relative to naïve neutrophils may be attributed to two potential mechanisms. Firstly, it is possible that the TNFα-priming have increased the plasma membrane expression of one or both GPCRs. Secondly, TNFα priming may have initiated intracellular signals that prime the NADPH oxidase itself for increased activity in response to certain stimuli. However, the subcellular location of BLT_1_R and FFA2R in human neutrophils remains to be elucidated, which is a challenging aspect to ascertain due to the unfortunate but well-known fact that no commercially available receptor antibodies are sufficiently selective and/or potent [58]. Moreover, the intracellular signalling pathways that are responsible for the assembly and subsequent activation of the NADPH oxidase are not yet fully understood [30, 59, 60], which limits the ability to speculate on the priming mechanism by TNFα. Nevertheless, the results of this study, in conjunction with those of previous investigations into GPCR-mediated elevations in [Ca^2+^]_i_ and NADPH oxidase activation in human neutrophils, indicate that an increase in [Ca^2+^]_i_ resulting from GPCR agonist stimulation is an insufficient condition for the concomitant activation of the NADPH oxidase NADPH oxidase [30]. Furthermore, neither TNFα nor Cmp58 mediate, alone or in combination, any neutrophil NADPH oxidase activation [48, 57]. The demonstrated transactivation mechanism between BLT_1_R and FFA2R was significantly more prominent in TNFα-primed as compared to naïve neutrophils. It may therefore be hypothesised that such an activation *in vivo* would be restricted to inflamed tissues as the transmigration process and tissue inflammatory mediators render neutrophils primed in a manner that mimics *in vitro* TNFα-primed neutrophils [31, 49, 61-63]. Neutrophils are the first immune cells to arrive at an inflamed site in the tissue in response to a variety of chemottractants including LTB_4_ [64, 65] for which neutrophils themselves constitute as a major source [20, 22, 23]. The levels of LTB_4_ have been demonstrated to be increased in diverse inflammatory diseases, both systemically (e.g., rheumatoid arthritis, systemic sclerosis and different types of asthma) and locally (e.g., in pancreatic tumours, the lung tissue and bronchoalveolar lavage fluid of patients with tuberculosis, lung cancer and asthma) [17, 18, 66]. Hence, both primed neutrophils and heightened levels of LTB_4_ are present in inflamed tissues, but whether also endogenous positive allosteric FFA2R modulators are present remains to be investigated. Regardless, the pro-inflammatory feature of this novel transactivation mechanism between BLT_1_R and allosterically modulated FFA2Rs, whereby LTB_4_ is converted to a potent activator of the NADPH oxidase, should be considered in the ongoing search for ligands or reagents targeting BLT_1_R and/or LTB_4_ for the treatment of inflammatory diseases [9, 16, 17, 53, 67-69]. Our results collectively indicate that the endogenous inflammatory mediator LTB_4_ induces receptor transactivation signals that are potentially identical to intracellular crosstalk signals generated by several other human neutrophil-expressed GPCRs. These signals either directly or indirectly activate allosterically modulated FFA2R which subsequently leads to signals that activate the ROS-producing neutrophil NADPH oxidase. This type of receptor transactivation, which involves one GPCR (i.e., BLT_1_R) that is activated by LTB_4_, and another allosterically modulated GPCR (i.e., FFA2R), represents a new regulatory mechanism that controls receptor activation in neutrophils from the cytosolic of the plasma membrane.

## CRediT authorship contribution statement

**Yanling Wu**: Conceptualization, Methodology, Formal analysis, Investigation, Visualization. **Claes Dahlgren**: Conceptualization, Methodology, Validation, Writing – Review and Editing, Supervision. **Huamei Forsman**: Conceptualization, Methodology, Validation, Investigation, Writing – Review and Editing, Supervision, Funding acquisition. **Martina Sundqvist**: Conceptualization, Methodology, Validation, Formal analysis, Investigation, Writing – Original Draft, Visualization, Supervision, Funding acquisition.

## Abbreviations

AA: arachidonic acid
AM: acetoxymethyl ester
AUC: area under the curve
BLTRs: leukotriene B_4_ receptors
BSA: bovine serum albumin
CATPB: FFA2R antagonist - (S)-3-[2-(3-chlorophenyl) acetamido]-4-[4-(trifluoromethyl)phenyl]butanoic acid
Cmp58: FFA2R allosteric modulator - ((*S*)-2-(4-chlorophenyl)-3,3-dimethyl-*N*-(5-phenylthiazol-2-yl)butanamide
EC_50_: half maximal effective concentration
FFA2R: free fatty acid receptor 2
fMLF: *N*-formyl-methionyl-leucyl-phenylalanine
FPRs: formyl peptide receptors
GM-CSF: granulocyte-macrophage colony-stimulating factor
GPCRs: G protein-coupled receptors
HRP: horseradish peroxidase
IP_3_: inositol-l,4,5-*tris*-phosphate
KRG: Krebs-Ringer glucose buffer
LPS: lipopolysaccharide
LTB_4_: Leukotriene B_4_
O_2_^-^: superoxide anion
PAFR: platelet activating factor receptor
PAM: positive allosteric modulator
PIP_2_: phosphatidyl-inositol-4,5-*bis*-phosphate
PLC: phospholipase C
ROS: reactive oxygen species
SCFAs: short chain fatty acids
TNFα: tumour necrosis factor α
U-75302: LTB_4_ receptor antagonist targeting BLT_1_R
YM-254890: Gα_q_ inhibitor
[Ca^2+^]_i_: intracellular concentration of free calcium ions (Ca^2+^)

## Acknowledgement

The work was financed by grants from the Swedish state under the agreement between the Swedish government and the county councils, the ALF-agreement (ALFGBG 78150), the Swedish Medical Research Council (2018-02848 and 2022-00624), the Åke Wiberg Foundation (M21-0025 and M23-0193), the Sahlgrenska International Starting Grant (GU2021/1070), the King Gustaf the V 80-year foundation (FAI-2021-0804 and FAI-2022-0873), the Rune and Ulla Almlövs Foundation (2023-418), the Mary von Sydow foundation (4723), the Magnus Bergwall foundation (2023-875), the Swedish Rheumatism Association (R-995669 and R-995361), the Health & Medical Care Committee of the Region Västra Götaland (VGFOUREG-979715 and VGFOUREG-995348), the Wilhelm and Martina Lundgren Science Fund (2024-SA-4605) and the Ingabritt and Arne Lundberg foundation. The sponsors did not have any role in any part of the study. The authors thank Linda Bergqvist for technical assistance regarding experiments.

## References

[1] M. Metzemaekers, B. Malengier-Devlies, M. Gouwy, L. De Somer, F.Q. Cunha, G. Opdenakker, P. Proost, Fast and furious: The neutrophil and its armamentarium in health and disease, Med Res Rev, 43 (2023) 1537–1606.

[2] S.B. Gacasan, D.L. Baker, A.L. Parrill, G protein-coupled receptors: the evolution of structural insight, AIMS Biophys, 4 (2017) 491–527.

[3] K.L. Pierce, R.T. Premont, R.J. Lefkowitz, Seven-transmembrane receptors, Nat Rev Mol Cell Biol, 3 (2002) 639–650.

[4] A.S. Hauser, M.M. Attwood, M. Rask-Andersen, H.B. Schiöth, D.E. Gloriam, Trends in GPCR drug discovery: new agents, targets and indications, Nat Rev Drug Discov, 16 (2017) 829–842.

[5] D. Kamato, P. Mitra, F. Davis, N. Osman, R. Chaplin, P.J. Cabot, R. Afroz, W. Thomas, W. Zheng, H. Kaur, M. Brimble, P.J. Little, Ga(q) proteins: molecular pharmacology and therapeutic potential, Cell Mol Life Sci, 74 (2017) 1379–1390.

[6] K. Sriram, P.A. Insel, G Protein-Coupled Receptors as Targets for Approved Drugs: How Many Targets and How Many Drugs?, Mol Pharmacol, 93 (2018) 251–258.

[7] C. Dahlgren, S. Lind, J. Mårtensson, L. Björkman, Y. Wu, M. Sundqvist, H. Forsman, G protein coupled pattern recognition receptors expressed in neutrophils: Recognition, activation/modulation, signaling and receptor regulated functions, Immunol Rev, 314 (2023) 69–92.

[8] A. Luginina, A. Gusach, E. Lyapina, P. Khorn, N. Safronova, M. Shevtsov, D. Dmitirieva, D. Dashevskii, T. Kotova, E. Smirnova, V. Borshchevskiy, V. Cherezov, A. Mishin, Structural diversity of leukotriene G-protein coupled receptors, J Biol Chem, 299 (2023) 105247.

[9] T. Yokomizo, T. Shimizu, The leukotriene B(4) receptors BLT1 and BLT2 as potential therapeutic targets, Immunol Rev, 317 (2023) 30–41.

[10] T. Shimizu, Lipid mediators in health and disease: enzymes and receptors as therapeutic targets for the regulation of immunity and inflammation, Annu Rev Pharmacol Toxicol, 49 (2009) 123–150.

[11] M. Bäck, W.S. Powell, S.E. Dahlén, J.M. Drazen, J.F. Evans, C.N. Serhan, T. Shimizu, T. Yokomizo, G.E. Rovati, Update on leukotriene, lipoxin and oxoeicosanoid receptors: IUPHAR Review 7, Br J Pharmacol, 171 (2014) 3551–3574.

[12] M. Nakamura, T. Shimizu, Recent advances in function and structure of two leukotriene B(4) receptors: BLT1 and BLT2, Biochem Pharmacol, 203 (2022) 115178.

[13] K. Saeki, T. Yokomizo, Identification, signaling, and functions of LTB(4) receptors, Semin Immunol, 33 (2017) 30–36.

[14] A.M. Tager, A.D. Luster, BLT1 and BLT2: the leukotriene B(4) receptors, Prostaglandins Leukot Essent Fatty Acids, 69 (2003) 123–134.

[15] A.W. Ford-Hutchinson, M.A. Bray, M.V. Doig, M.E. Shipley, M.J. Smith, Leukotriene B, a potent chemokinetic and aggregating substance released from polymorphonuclear leukocytes, Nature, 286 (1980) 264–265.

[16] F. Sasaki, T. Yokomizo, The leukotriene receptors as therapeutic targets of inflammatory diseases, Int Immunol, 31 (2019) 607–615.

[17] R. He, Y. Chen, Q. Cai, The role of the LTB4-BLT1 axis in health and disease, Pharmacol Res, 158 (2020) 104857.

[18] A. Jo-Watanabe, T. Okuno, T. Yokomizo, The Role of Leukotrienes as Potential Therapeutic Targets in Allergic Disorders, Int J Mol Sci, 20 (2019).

[19] S.W. Crooks, R.A. Stockley, Leukotriene B4, Int J Biochem Cell Biol, 30 (1998) 173–178.

[20] K. Kienle, T. Lämmermann, Neutrophil swarming: an essential process of the neutrophil tissue response, Immunol Rev, 273 (2016) 76–93.

[21] M. Peters-Golden, W.R. Henderson, Jr., Leukotrienes, N Engl J Med, 357 (2007) 1841–1854.

[22] T. Lämmermann, In the eye of the neutrophil swarm-navigation signals that bring neutrophils together in inflamed and infected tissues, J Leukoc Biol, 100 (2016) 55–63.

[23] T. Németh, A. Mócsai, Feedback Amplification of Neutrophil Function, Trends Immunol, 37 (2016) 412–424.

[24] G. Lärfars, F. Lantoine, M.A. Devynck, J. Palmblad, H. Gyllenhammar, Activation of nitric oxide release and oxidative metabolism by leukotrienes B4, C4, and D4 in human polymorphonuclear leukocytes, Blood, 93 (1999) 1399-1405.

[25] J. Palmblad, H. Gyllenhammar, J.A. Lindgren, C.L. Malmsten, Effects of leukotrienes and f-Met-Leu-Phe on oxidative metabolism of neutrophils and eosinophils, J Immunol, 132 (1984) 3041–3045.

[26] M.P. Wymann, V. von Tscharner, D.A. Deranleau, M. Baggiolini, The onset of the respiratory burst in human neutrophils. Real-time studies of H2O2 formation reveal a rapid agonist-induced transduction process, J Biol Chem, 262 (1987) 12048-12053.

[27] R.H. Weisbart, L. Kwan, D.W. Golde, J.C. Gasson, Human GM-CSF primes neutrophils for enhanced oxidative metabolism in response to the major physiological chemoattractants, Blood, 69 (1987) 18–21.

[28] H. Gyllenhammar, Mechanisms for luminol-augmented chemiluminescence from neutrophils induced by leukotriene B4 and N-formyl-methionyl-leucyl-phenylalanine, Photochem Photobiol, 49 (1989) 217–223.

[29] H. Gyllenhammar, Correlation between neutrophil superoxide formation, luminol-augmented chemiluminescence and intracellular Ca2+ levels upon stimulation with leukotriene B4, formylpeptide and phorbolester, Scand J Clin Lab Invest, 49 (1989) 317–322.

[30] C. Dahlgren, H. Forsman, M. Sundqvist, L. Björkman, J. Mårtensson, Signaling by Neutrophil G Protein-Coupled Receptors that Regulate the Release of Superoxide Anions, J Leukoc Biol, (2024).

[31] J.B. Cowland, N. Borregaard, Granulopoiesis and granules of human neutrophils, Immunol Rev, 273 (2016) 11–28.

[32] I. Miralda, S.M. Uriarte, K.R. McLeish, Multiple Phenotypic Changes Define Neutrophil Priming, Front Cell Infect Microbiol, 7 (2017) 217.

[33] K.L. Vogt, C. Summers, A.M. Condliffe, The clinical consequences of neutrophil priming, Curr Opin Hematol, 26 (2019) 22–27.

[34] I. Niemietz, A.T. Moraes, M. Sundqvist, K.L. Brown, Hyaluronan primes the oxidative burst in human neutrophils, J Leukoc Biol, 108 (2020) 705–713.

[35] M. Sundqvist, A. Holdfeldt, S.C. Wright, T.C. Møller, E. Siaw, K. Jennbacken, H. Franzyk, M. Bouvier, C. Dahlgren, H. Forsman, Barbadin selectively modulates FPR2-mediated neutrophil functions independent of receptor endocytosis, Biochim Biophys Acta Mol Cell Res, 1867 (2020) 118849.

[36] C. Dahlgren, A. Holdfeldt, S. Lind, J. Mårtensson, M. Gabl, L. Björkman, M. Sundqvist, H. Forsman, Neutrophil Signaling That Challenges Dogmata of G Protein-Coupled Receptor Regulated Functions, ACS Pharmacol Transl Sci, 3 (2020) 203–220.

[37] S. Lind, K.L. Granberg, H. Forsman, C. Dahlgren, The allosterically modulated FFAR2 is transactivated by signals generated by other neutrophil GPCRs, PLoS One, 18 (2023) e0268363.

[38] S. Lind, A. Holdfeldt, J. Mårtensson, M. Sundqvist, L. Björkman, H. Forsman, C. Dahlgren, Functional selective ATP receptor signaling controlled by the free fatty acid receptor 2 through a novel allosteric modulation mechanism, Faseb j, 33 (2019) 6887–6903.

[39] A. Böyum, Isolation of mononuclear cells and granulocytes from human blood. Isolation of monuclear cells by one centrifugation, and of granulocytes by combining centrifugation and sedimentation at 1 g, Scand J Clin Lab Invest Suppl, 97 (1968) 77-89.

[40] A. Bøyum, Isolation of lymphocytes, granulocytes and macrophages, Scand J Immunol, Suppl 5 (1976) 9–15.

[41] L. Björkman, H. Forsman, L. Bergqvist, C. Dahlgren, M. Sundqvist, Larixol is not an inhibitor of Gα(i) containing G proteins and lacks effect on signaling mediated by human neutrophil expressed formyl peptide receptors, Biochem Pharmacol, 220 (2024) 115995.

[42] J. Bylund, H. Björnsdottir, M. Sundqvist, A. Karlsson, C. Dahlgren, Measurement of respiratory burst products, released or retained, during activation of professional phagocytes, Methods Mol Biol, 1124 (2014) 321–338.

[43] C. Dahlgren, H. Björnsdottir, M. Sundqvist, K. Christenson, J. Bylund, Measurement of Respiratory Burst Products, Released or Retained, During Activation of Professional Phagocytes, Methods Mol Biol, 2087 (2020) 301–324.

[44] K. Futosi, S. Fodor, A. Mócsai, Reprint of Neutrophil cell surface receptors and their intracellular signal transduction pathways, Int Immunopharmacol, 17 (2013) 1185–1197.

[45] G.M. Bokoch, Chemoattractant signaling and leukocyte activation, Blood, 86 (1995) 1649–1660.

[46] Y. Wang, C.L. Zhu, P. Li, Q. Liu, H.R. Li, C.M. Yu, X.M. Deng, J.F. Wang, The role of G protein-coupled receptor in neutrophil dysfunction during sepsis-induced acute respiratory distress syndrome, Front Immunol, 14 (2023) 1112196.

[47] H. Forsman, Y. Wu, J. Mårtensson, L. Björkman, K.L. Granberg, C. Dahlgren, M. Sundqvist, AZ2158 is a more potent formyl peptide receptor 1 inhibitor than the commonly used peptide antagonists in abolishing neutrophil chemotaxis, Biochem Pharmacol, 211 (2023) 115529.

[48] J. Mårtensson, A. Holdfeldt, M. Sundqvist, M. Gabl, T.P. Kenakin, L. Björkman, H. Forsman, C. Dahlgren, Neutrophil priming that turns natural FFA2R agonists into potent activators of the superoxide generating NADPH-oxidase, J Leukoc Biol, 104 (2018) 1117–1132.

[49] M. Sundqvist, K. Christenson, A. Holdfeldt, M. Gabl, J. Mårtensson, L. Björkman, R. Dieckmann, C. Dahlgren, H. Forsman, Similarities and differences between the responses induced in human phagocytes through activation of the medium chain fatty acid receptor GPR84 and the short chain fatty acid receptor FFA2R, Biochim Biophys Acta Mol Cell Res, 1865 (2018) 695–708.

[50] S. Lind, Y. Wu, M. Sundqvist, H. Forsman, C. Dahlgren, An increase in the cytosolic concentration of free calcium ions activates the neutrophil NADPH-oxidase provided that the free fatty acid receptor 2 has been allosterically modulated, Cell Signal, 107 (2023) 110687.

[51] A. Bock, M. Bermudez, Allosteric coupling and biased agonism in G protein-coupled receptors, Febs j, 288 (2021) 2513–2528.

[52] D. Wootten, A. Christopoulos, P.M. Sexton, Emerging paradigms in GPCR allostery: implications for drug discovery, Nat Rev Drug Discov, 12 (2013) 630–644.

[53] M. Grundmann, E. Bender, J. Schamberger, F. Eitner, Pharmacology of Free Fatty Acid Receptors and Their Allosteric Modulators, Int J Mol Sci, 22 (2021).

[54] T.S. Peterson, J.M. Camden, Y. Wang, C.I. Seye, W.G. Wood, G.Y. Sun, L. Erb, M.J. Petris, G.A. Weisman, P2Y2 nucleotide receptor-mediated responses in brain cells, Mol Neurobiol, 41 (2010) 356–366.

[55] H. Jin, Y.S. Ko, S.P. Yun, S.W. Park, H.J. Kim, P2Y(2)R-mediated transactivation of VEGFR2 through Src phosphorylation is associated with ESM-1 overexpression in radiotherapy-resistant-triple negative breast cancer cells, Int J Oncol, 62 (2023).

[56] L.T. Woods, K.J. Jasmer, K. Muñoz Forti, V.C. Shanbhag, J.M. Camden, L. Erb, M.J. Petris, G.A. Weisman, P2Y(2) receptors mediate nucleotide-induced EGFR phosphorylation and stimulate proliferation and tumorigenesis of head and neck squamous cell carcinoma cell lines, Oral Oncol, 109 (2020) 104808.

[57] L. Björkman, J. Mårtensson, M. Winther, M. Gabl, A. Holdfeldt, M. Uhrbom, J. Bylund, A. Højgaard Hansen, S.K. Pandey, T. Ulven, H. Forsman, C. Dahlgren, The Neutrophil Response Induced by an Agonist for Free Fatty Acid Receptor 2 (GPR43) Is Primed by Tumor Necrosis Factor Alpha and by Receptor Uncoupling from the Cytoskeleton but Attenuated by Tissue Recruitment, Mol Cell Biol, 36 (2016) 2583–2595.

[58] R. Ayoubi, J. Ryan, M.S. Biddle, W. Alshafie, M. Fotouhi, S.G. Bolivar, V. Ruiz Moleon, P. Eckmann, D. Worrall, I. McDowell, K. Southern, W. Reintsch, T.M. Durcan, C. Brown, A. Bandrowski, H. Virk, A.M. Edwards, P. McPherson, C. Laflamme, Scaling of an antibody validation procedure enables quantification of antibody performance in major research applications, Elife, 12 (2023).

[59] W.M. Nauseef, Assembly of the phagocyte NADPH oxidase, Histochem Cell Biol, 122 (2004) 277–291.

[60] W.M. Nauseef, The phagocyte NOX2 NADPH oxidase in microbial killing and cell signaling, Curr Opin Immunol, 60 (2019) 130–140.

[61] A. Karlsson, P. Follin, H. Leffler, C. Dahlgren, Galectin-3 activates the NADPH-oxidase in exudated but not peripheral blood neutrophils, Blood, 91 (1998) 3430–3438.

[62] H. Sengeløv, P. Follin, L. Kjeldsen, K. Lollike, C. Dahlgren, N. Borregaard, Mobilization of granules and secretory vesicles during in vivo exudation of human neutrophils, J Immunol, 154 (1995) 4157–4165.

[63] M. Sundqvist, A. Welin, J. Elmwall, V. Osla, U.J. Nilsson, H. Leffler, J. Bylund, A. Karlsson, Galectin-3 type-C self-association on neutrophil surfaces; The carbohydrate recognition domain regulates cell function, J Leukoc Biol, 103 (2018) 341–353.

[64] M. Metzemaekers, M. Gouwy, P. Proost, Neutrophil chemoattractant receptors in health and disease: double-edged swords, Cell Mol Immunol, 17 (2020) 433–450.

[65] W.M. Nauseef, N. Borregaard, Neutrophils at work, Nat Immunol, 15 (2014) 602–611.

[66] G.Y. Moore, G.P. Pidgeon, Cross-Talk between Cancer Cells and the Tumour Microenvironment: The Role of the 5-Lipoxygenase Pathway, Int J Mol Sci, 18 (2017).

[67] L. Bhatt, K. Roinestad, T. Van, E.B. Springman, Recent advances in clinical development of leukotriene B4 pathway drugs, Semin Immunol, 33 (2017) 65–73.

[68] S.L. Brandt, C.H. Serezani, Too much of a good thing: How modulating LTB(4) actions restore host defense in homeostasis or disease, Semin Immunol, 33 (2017) 37–43.

[69] I. Kimura, A. Ichimura, R. Ohue-Kitano, M. Igarashi, Free Fatty Acid Receptors in Health and Disease, Physiol Rev, 100 (2020) 171–210.

